# On the impact of reference selection on variant calling in phylogenomics: How to avoid systematic error in target-enrichment studies of non-model organisms

**DOI:** 10.1101/2025.03.23.644254

**Authors:** Guoyi Zhang, Gerasimos Cassis, Frank Köhler

## Abstract

Target-enrichment methods are widely used in phylogenomic studies aiming to reconstruct the relationships among diverse groups of organisms. When building phylogenomic datasets from short sequencer reads, software pipelines need to accurately align short sequencer reads to the target and identify the genetic variation in homologous DNA sequences. Variant calling, a crucial step in this process, is known to be error-prone in a phylogenetic analysis framework as it requires a conspecific reference as a template. To avoid problems relating to reference choice, existing target-enrichment pipelines often skip variant calling entirely using work-arounds. This study investigates the effect of reference genome selection on the reliability of variant calling and assesses the consequences of reference genome choice on phylogenomic analyses downstream. Employing an empirical exon capture sequence dataset, we examine how variant detection is influenced by reference genome choice. We reconstructed and analysed phylogenomic sequence datasets from short sequencer reads, which includes steps of quality control, read mapping, variant calling, multiple sequence alignment and phylogenetic analysis. We assembled the same exon capture dataset by using four different references for variant calling. These references were (1) a distantly related species, (2) a single ingroup sample, (3) a composite of all ingroup samples, and (4) a self-derived reference for each sample. We found that reference choice significantly impacted the variant detection and that these differences in variant detection influenced the phylogenetic reconstructions downstream. Comparing multiple sequence alignments and phylogenetic trees produced using different references revealed that employing a sample-specific self-reference for variant calling maximizes the accuracy of the variant detection process. Based on this finding, we recommend incorporating self-referenced variant calling in the assembly of phylogenomic datasets to ensure robustness and reproducibility, especially for datasets characterized by low read coverage.

## Introduction

Next-generation sequencing technologies have revolutionized biodiversity research by enabling the generation of vast amounts of genomic data, thereby opening unprecedented opportunities to investigate the genetic basis of organismal diversity (Dijk et al. 2014). High-throughput sequencing platforms produce millions of short DNA fragments (reads) that cannot be directly used for phylogenetic analysis. To process these data, phylogenomic studies employ bioinformatic pipelines – automated workflows that integrate multiple specialized software tools to transform raw sequence reads into datasets of homologous sequences suitable for evolutionary inference. This transformation typically involves three stages: (i) data acquisition and preprocessing, (ii) gene and ortholog identification, and (iii) sequence alignment (Roy et al. 2018).

The first stage includes quality assessment, sequence filtering, and the alignment of reads to contiguous sequences (contigs) by stitching together overlapping reads based on shared sequence patterns with a reference genome (Langmead and Salzberg 2012). In this process, the reference genome provides a template for identifying, assembling, and aligning orthologous loci. In addition, it plays a vital role in the detection of sequence variants (Li et al. 2009; DePristo et al. 2011; Link et al. 2017; Lefouili and Nam 2022).

The short-read alignment and the detection of sequence variants, a process called variant calling, are two processes. Aligners, such as bwa (Li & Durbin 2009) and bowtie2 (Langmead & Salzberg, 2012), map reads against the reference (Langmead & Salzberg, 2012), and variant callers, such as BCFtools (Danecek et al. 2021) and GATK (DePristo et al. 2011), identify nucleotide differences among these reads with respect to the reference (DePristo et al. 2011; Link et al. 2017; Danecek et al. 2021). Variant callers distinguish true nucleotide differences from sequencing errors by relying on several key assumptions about how sequencing data behave (Li et al. 2009; Li 2011; Link et al. 2017; Danecek et al. 2021). For example, they assume that sequencing errors occur randomly and at low frequency, whereas true variants are consistently supported by multiple high-quality reads. Correct read mapping and adequate coverage are also assumed, as misalignments or low read depth can mimic false variants (Li 2011; Nevado et al. 2014; Yun et al. 2020; Deng et al. 2022). Variant callers further expect that base quality scores accurately reflect error probabilities and that allele frequencies follow biologically plausible ratios (Li 2011; Yun et al. 2020; Rick et al. 2024). By integrating these assumptions within a probabilistic framework calibrated to the sequencer’s empirical error profile, variant callers can infer genuine polymorphisms while minimizing false calls (Olson et al. 2015). The output of the short-read alignment and variant process are consensus sequence of the target contigs to move on to the next step of the data transformation mentioned above.

While now widely used in population genetics and phylogenomics, variant calling was originally developed for detecting heterozygosity in diploid human genomes (Li et al. 2009; Link et al. 2017; Lefouili and Nam 2022; DePristo et al. 2011; Li 2011). Therefore, to function accurately, variant callers require a reference genome that is closely related to (i.e., conspecific with) the analyzed samples. That means that in phylogenomic studies of non-model organisms difficulties may arise from the unavailability of conspecific reference genomes (e.g., Duchen and Salamin, 2021).

Duchen and Salamin (2021) cautioned that variant callers can substantially underestimate the number of variants when applied at a phylogenetic rather than population-genetic scale, and they showed that some callers perform better than others under these conditions. More recently, Rick et al. (2024) examined the influence of reference-genome choice in phylogenomic analyses in greater depth. Using both simulated and empirical datasets, they demonstrated that thresholds for filtering minor alleles at heterozygous sites during short-read alignment can alter phylogenomic outcomes. Importantly, they also showed that reference-genome selection can introduce biases that propagate through downstream analyses as variant calling becomes increasingly inaccurate. Such inaccuracies ultimately compromise the reliability and reproducibility of phylogenomic reconstructions. Although Rick et al. (2024) concluded that reference-genome choice has significant implications for variant-calling accuracy and subsequent analyses, their study did not offer practical strategies for fully mitigating these problems.

Without a practical solution, several pipelines were designed to assemble target-enrichment datasets from phylogenetically divergent samples by avoiding variant calling altogether. Instead, these pipelines rely on *de novo* assembly of contigs and sequence matching against references using BLAST [e.g., HybPiper (Johnson et al. 2016), PHYLUCE (Faircloth 2016) and Assexon (Yuan et al. 2019)]. Hugall et al. (2016) mapped reads against a so-called ‘superreference’ made up of multiple references. However, variant calling enhances the accuracy and completeness of the final contigs that are used to build the final multiple sequence alignment (MSA).

Many phylogenomic studies face the problem that closely related reference genomes for the organisms under study are not available. To address the problem of variant calling in these cases, we use this pipeline to assess the influence of different reference genomes on variant calling. To do this, we assembled exon capture sequences using different reference exomes for variant calling (Fig. 1). These references were (1) a distantly related species (for run 1), (2) a single ingroup sample (for runs 2-23), (3) a composite of all ingroup samples (for run 24), and (4) a self-derived reference for each sample (for run 25). We calculated pairwise distances to quantify the variant differences between contigs assembled using different references. We compared phylogenies based on MSAs that were reconstructed using different reference genomes to assess their impact on downstream phylogenomic analyses. Finally, we checked short-read alignment and the corresponding called variants by eye.

**Fig. 1.**
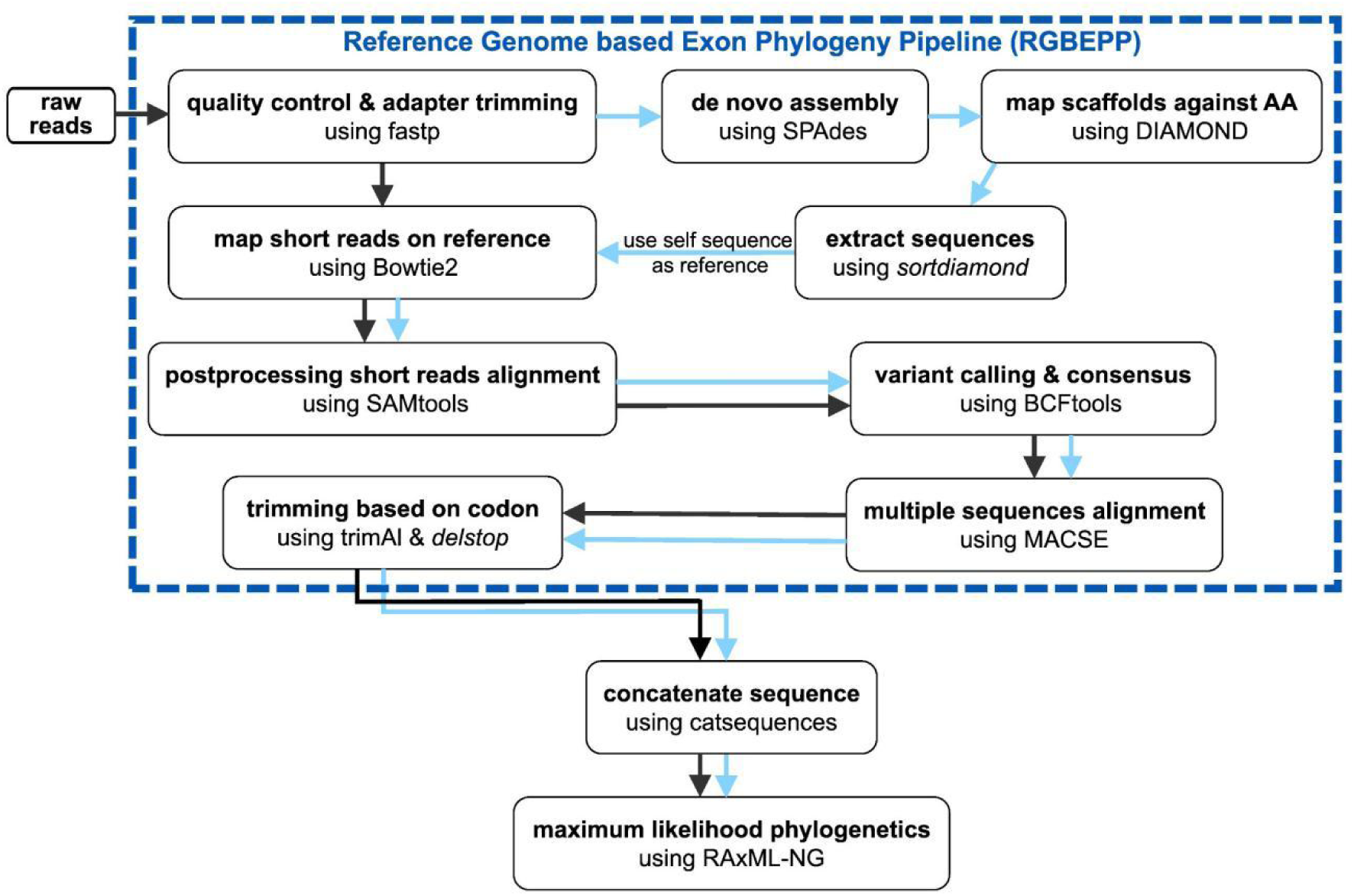
Overview of the two workflows RGBEPP (blue arrows, for run 25) and RGBEPP_refmix (black arrows, for runs 1-24) used herein. New tools provided in this study are in Italics.

Based on our observations, we provide practical recommendations to maximize the accuracy of variant discovery in phylogenomic datasets. Specifically, we propose a novel assemblysolution for target-enrichoment phylogenomic dataset, which limits inaccuracies in variant calling based on reference choice.

## Results

### Data Characteristics

The multiple sequence alignment (MSA) with *Lottia gigantea* as a reference had a length of 144,831 bp. Sequence alignments using other references varied from that length insignificantly depending on the completeness and quality of the references used.

### Genetic distances among MSAs

We detected substantial differences in the calculated pairwise distances among the MSAs produced under varying choices of reference exomes, which are visualized in heatmaps (Fig. 2). These heatmaps reveal that the lowest interspecific distances were generally observed in the MSA produced for the outgroup variant calling reference (run 1). In this case, the interspecific pairwise distances between aligned partial exomes were 0. These low distances mean that almost no variants were detected. The MSAs produced by using ingroup samples or combined ingroup as references for variant calling (runs 2-24) revealed overall higher interspecific distances between aligned exons (single from 0.004 to 0.041; combined from 0.012 to 0.037). Consequently, a larger proportion of variants were called. The largest distances were observed in the MSA produced under self-referenced variant calling (run 25: 0.020 to 0.108) indicating that this referencing method scored the most differences (Fig. 2, S1).

**Fig. 2.**
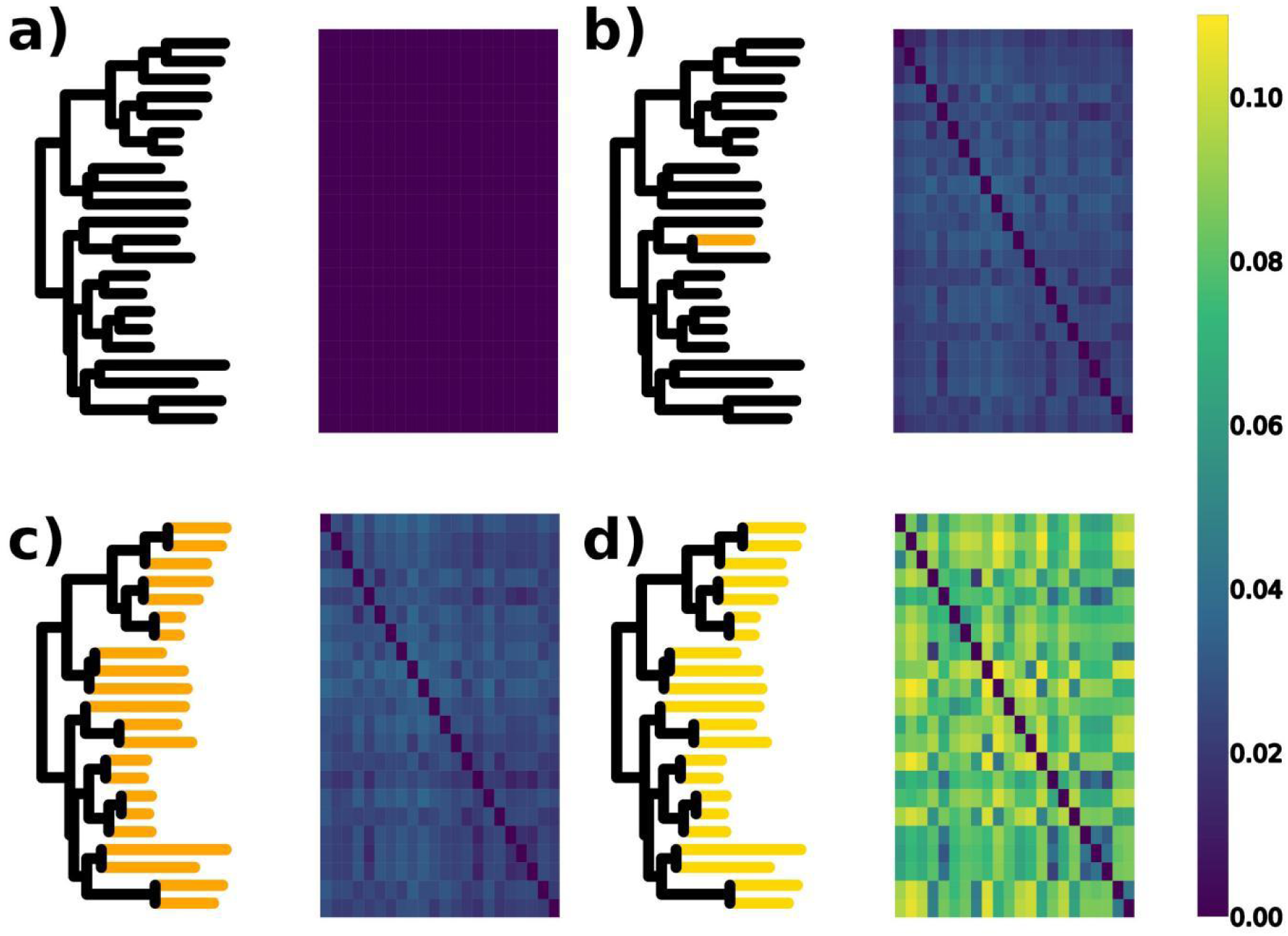
Heatmaps of pairwise genetic distances among the partial exomes in each of the MSAs built using different reference exomes. a) non-ingroup reference (run 1); b) individual ingroup references (runs 2), referenced taxon highlighted in corresponding tree; c) combined ingroup reference (run 24); d) sample-specific self-reference (run 25).

### Genetic distances among exomes of same samples

The p-distances between partial exomes of the same samples revealed differences in the proportion of called variants under varying references. Plotting these distances in a MDS diagram revealed two well-separated clouds: One cloud represents all assemblies produced for the outgroup reference (run 1) while the second cloud contains all other assemblies (runs 2-25) (Figs 3, S2). The significant separation of these two clouds indicates substantial differences in the proportion of called variants.

**Fig. 3.**
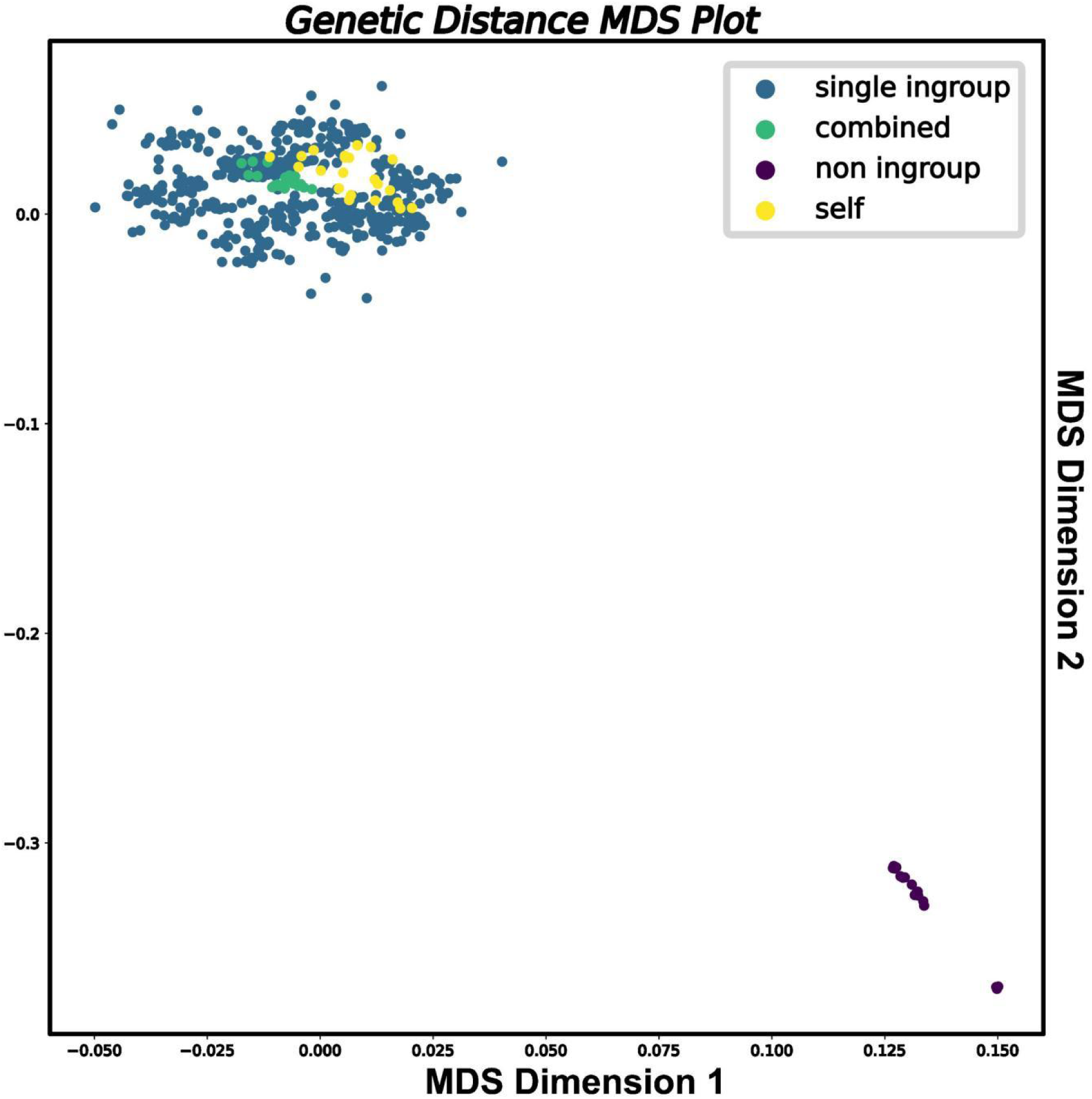
Scatter plot of multidimensional scaling (MDS) summarizing p-distances among ingroup taxa for each MSA. Colours indicate reference used for variant calling.

The spread of the cloud for various ingroup references (runs 2-25) shows that there are differences in variant detection also, but these differences were less substantial in comparison with the differences to run 1.

### Comparisons of tree topologies

Modelfinder identified the general time-reversible model of sequence evolution with gamma distributed rates (GTR+I+G) (Tavaré, 1986) as the best model for all phylogenetic analyses but one. For the dataset produced with the outgroup representative as a reference, it identified the F81 model (Felsenstein 1981).

The differences in the proportion of called variants did not only figure in varied pairwise genetic distances among and between the assembled partial exomes as shown above. More importantly, these differences also affected the topologies of the Maximum Likelihood trees reconstructed for the different MSAs. Using relative Robinson-Foulds (RF) distances to quantify the observed topological differences among the different phylogenetic trees (Fig. 4), we found that the tree topologies based on single ingroup references (runs 2-23) varied from one another as well as from the topologies of the other trees. By contrast, the trees produced under the self-referenced (run 25) and combined ingroup (run 24) methods were rather similar to one another. The tree reconstructed for the MSA generated with *Lottia gigantea* as a reference (run 1) was least similar with any other tree.

**Fig. 4.**
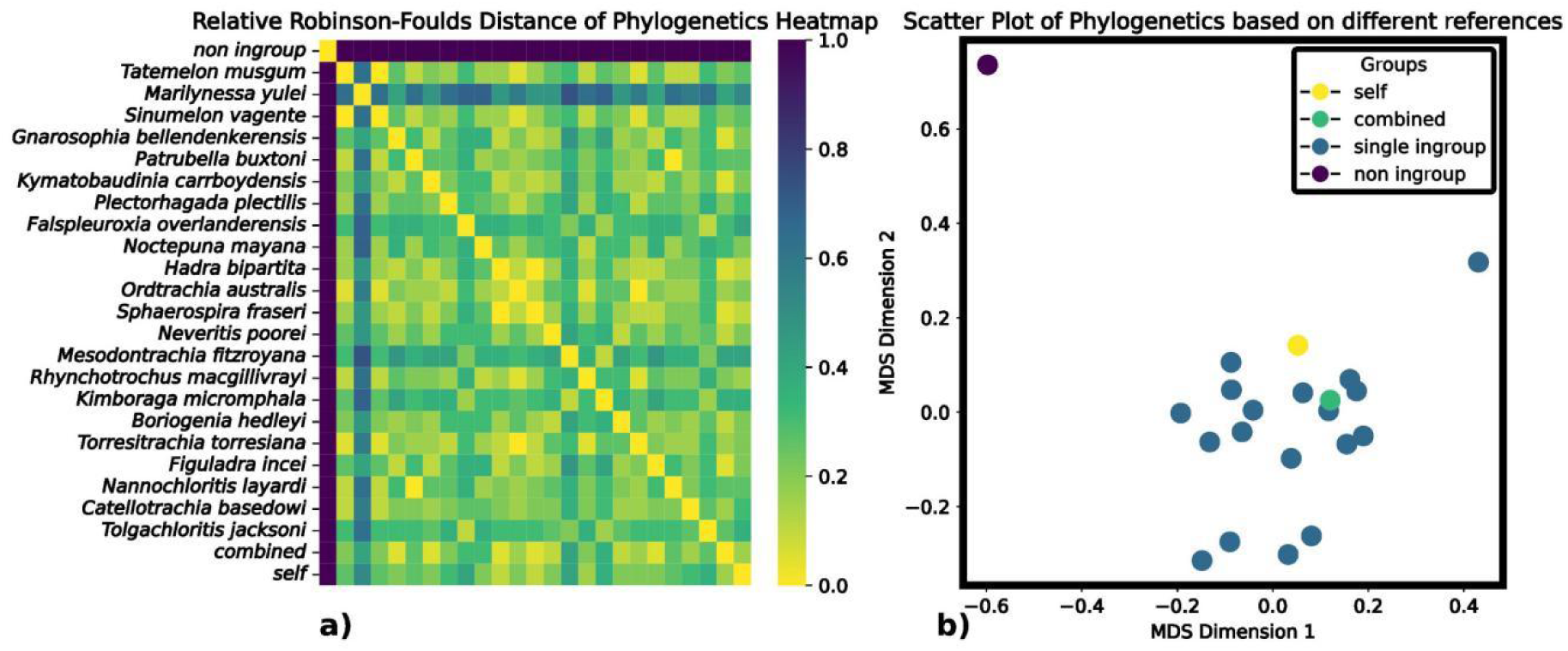
a) Heatmap of relative Robinson-Foulds (RF) distances between all trees based on MSAs built under different reference exome choices. b) Multidimensional Scaling scatter plot of relative RF distances for these trees. Colours indicate reference used for variant calling.

## Discussion

### The impact of reference choice on phylogenomic analyses

Phylogenomic studies, including target enrichment studies, usually require the use of reference sequences as templates for identifying, assembling, and aligning orthologous loci from millions of short sequencing reads. The critical role of reference genomes in variant calling workflows was highlighted in previous studies (Hwang et al. 2015; Cornish and Guda, 2015; Chen et al. 2019; Barbitoff et al. 2022). When comparing their efficacy in phylogenomic studies, Duchen and Salamin (2021) noted that the choice of variant caller software can significantly influence the results of phylogenomic analyses. Subsequently, Rick et al. (2024) demonstrated that both the parameters of the variant callers and the choice of the reference genome impact sequence assembly as well as the phylogenetic reconstructions downstream. They argued that if a reference is too divergent from the target samples, then reads may map incorrectly or fail to map altogether. Proposed strategies to mitigate the impact of reference choice include using genomes of a closely related species as a reference or building a composite reference from several taxa (Duchen and Salamin 2021, Rick et al. 2024).

Our trials of constructing MSAs using different references for variant calling aimed to test the adequacy of these two strategies in avoiding systematic error during the assembly of short reads. In addition, we compared the performance of a novel strategy proposed herein for aligning short reads against a sample-specific self-reference.

The design of our experiments is simple. We assembled short reads repeatedly from scratch whereby the only varying factor was the chosen reference or the way in which that reference was constructed. Then we calculated p-distances between (1) all sequences in each MSA and (2) between the same contig sequences assembled with different references to illustrate the variation in the number of detected variants. Any difference in p-distances can be attributed to a varying number of called variants. In addition, the p-distances provide a measure for the magnitude of the problem.

We found that using a phylogentically distant reference for variant calling in data from a closely related species significantly reduced the numbers of discovered variants overall.ll. Because of the extremely low number of called variants, the measured distances between the sequences in the assembled dataset were very low in this case, in effect close to zero. This observation suggests that ‘false negatives’, or uncalled variants, appear to be the main source of error in the variant calling process. This error decreases with a decreasing phylogenetic distance between the reference and the sample. This conclusion can be drawn from the larger numbers of variants called in runs 2-24 that used references that were phylogenetically closer to the templates. There are two main sources of error: First, when the reference is highly divergent from the sample (i.e., a non-ingroup reference), short reads may fail to align at all, leading to substantial data loss and an underestimation of true variation. Secondly, at intermediate levels of divergence between reference and sample (i.e., an ingroup reference), reads may align reasonably well, but variant calling is biased toward the reference allele, resulting in the under-calling of true variants, causing false negatives. The mechanisms of this error are illustrated in IGV plots of a short-read alignment depicted in Supplementary Figure 5. When using a single, closely related reference, multiple nucleotide differences between this reference and the mapped reads of the sample are detected. However, few of these detected differences are called as variants while most are not (Fig. 5a). As a result, the consensus sequence created by the variant caller differs from the reference in many fewer nucleotides than from the sequence of the mapped reads of the sample. The nucleotide differences between reference and mapped reads that are not called as variants are almost exclusively false negatives. When a composite reference is used, the number of observed nucleotide differences between the mapped reads of the sample and any of the references is minimized and nearly all observed differences are called as variants (Fig. 5b). However, which variants are called correctly is not transparent for the heterogenous nature of the composite references. In the case of self-referencing, very few nucleotide differences between the reference and the mapped reads are observed. In the present example, a single nucleotide difference, presumably representing a sequencing error, is rightly not called as a variant because of the very low frequency with which it occurs across the mapped reads (Fig. 5c). The described bias in variant calling can introduce spurious similarity across the sequences in the dataset through incorporating states from the reference into the alignment, which are false as they are not represented in the sample reads. Consequently, the number and distribution of detected variants depend strongly on the degree of divergence between reference and sample. The most reliable variant calling occurs when this divergence is minimal.

**Fig. 5.**
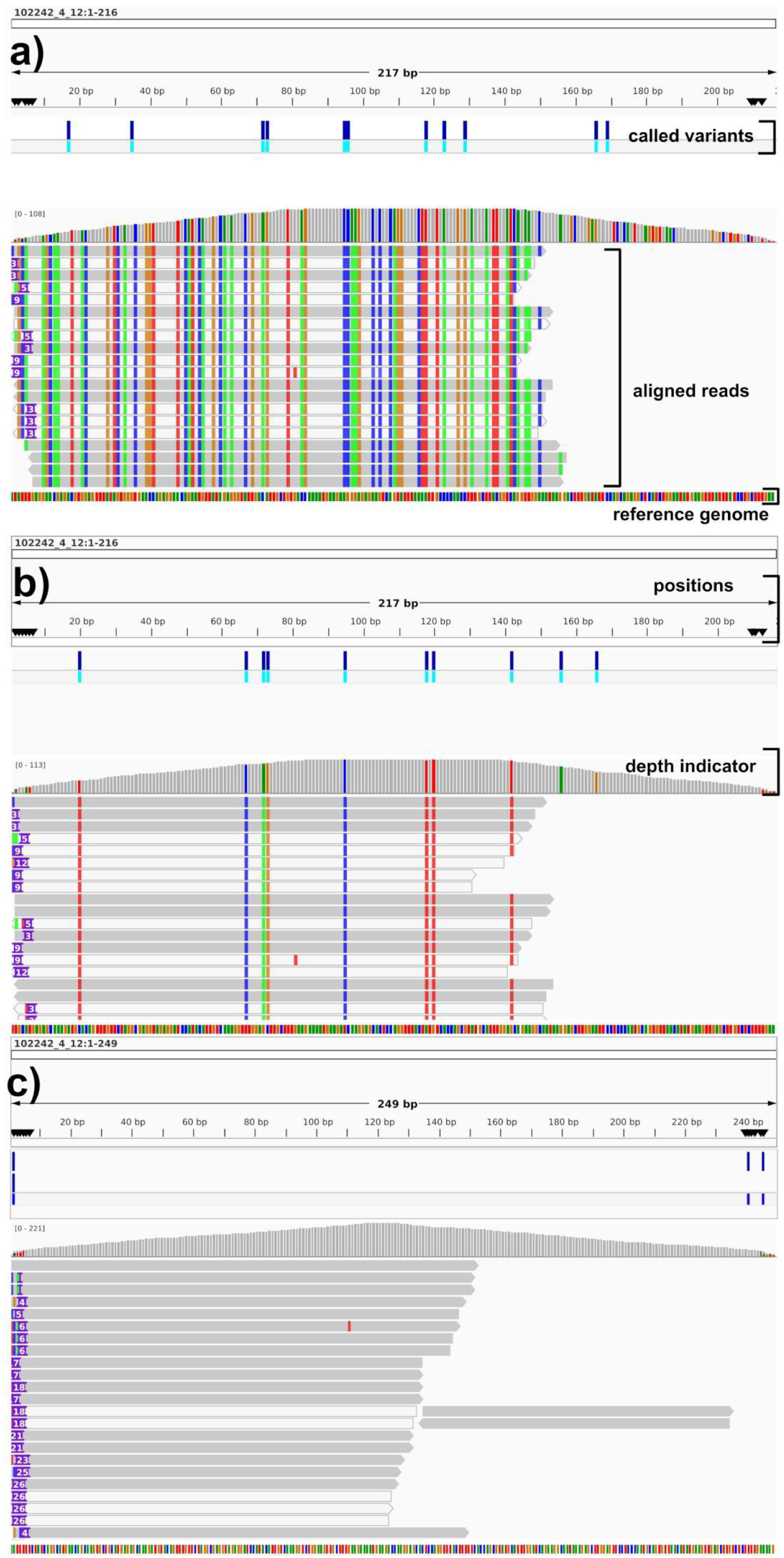
IGV screenshots of short read alignment with called variants. Horizontal lines indicate individual sequencer reads. Grey indicates invariant DNA sequences. Coloured columns indicate observed nucleotide differences among reads and reference genome. a) Short-read alignment using a single reference genome; b) Short-read alignment using a composite reference; c) Short-read alignment using a self-referencing.

Our observation that a phylogenetically distant reference is unsuitable for short-read alignment and variant calling is unsurprising given the studies of Duchen and Salamin (2021) and Rick et al. (2024). However, choosing references that are more closely related to the target group, as suggested as a remedy for this problem, does not fully remove all variant calling errors as we show here. We found that among the MSAs assembled using in-group references (runs 2-23), the proportions of detected variants varied stochastically and noticeably (Figs 2, 3). In a few cases, a large proportion of called variants even accumulated in a few samples (e.g., Figure S1e). Using a composite reference from several taxa (here all 22 ingroup samples, run 24) did not produce a MSA that contained noticeably more phylogenetic information than any of the MSAs produced for a single ingroup reference indicating that variant calling is still plagued by false negatives (run 24, Figure S1x). By contrast, we show that the p-distances among all samples are noticeably larger when self-referencing is used (run 25, Figure S1y). These overall larger genetic distances are testimony to an overall larger proportion of variants being called. Because false negatives are the main source of error, a larger number of called variants reduces that error. The largest number of variants, equaling the smallest error, is called when the genetic divergence between reference and sample is zero.

Employing relative Robinson-Foulds (RF) distances, we demonstrate that the above-mentioned variations in the numbers of called variants impact the topologies of phylogenetic trees. Plotting these distances (Fig. 4) shows that the tree based on the MSA assembled for a distantly related reference is significantly different from all other trees. In addition, this plot also shows that there are detectable topological differences among all trees based on MSAs assembled for in-group references (runs 2-25).

In summary, our observations corroborate the conclusions of previous studies that more distantly related reference genomes tend to yield fewer accurate variant calls and that these inaccuracies have impacted phylogenetic inferences significantly. Therefore, it is important that methods are implemented to minimize the error resulting from reference selection. We show that this error decreases with the phylogenetic distance between reference and sample sequences and conclude that ideally distance should be near zero (i.e., reference and sample are conspecific).

### Possible causes for differences in variant calling

In principle, the discrepancies between the MSAs built using varying references may result from two factors: the behavior of different variant callers, and the characteristics of the datasets. The suitability of widely used variant callers, such as GATK HaplotypeCaller (DePristo et al. 2011), FreeBayes (Garrison and Marth 2012) and BCFtools (Danecek et al. 2021), for phylogenomic studies was addressed previously by studies highlighting performance differences among these programs (e.g., Hwang et al. 2015; Lefouili and Nam 2022; Duchen and Salamin, 2021). However, these differences are marginal in comparison with the magnitude of the differences reported herein.

In a phylogenomic context, a major limitation of variant callers arises from the fact that these programs were originally developed for population genomic analyses rather than for multi-species phylogenomic datasets (DePristo et al. 2011; Li 2011). Variant calling algorithms assume that sequencing reads are mapped against a reference genome from the same species. When the reference genome is phylogenetically divergent from the sample, then pre-existing nucleotide differences between reference and sample can mislead the variant calling process. The underlying mechanisms are complex: reads fail to align in highly divergent regions, true variants may not be detected, and allele frequencies are distorted (Rick et al. 2024). As a result, the variant sets produced under these conditions violate assumptions such as the Hardy–Weinberg equilibrium (Yun et al. 2020) and introduce systematic error that compromises downstream analyses.

While the impact of computational tools and parameters used for variant calling on genomic analyses has been widely discussed (Hwang et al. 2015; Cornish and Guda 2015; Chen et al. 2019; Barbitoff et al. 2022), little attention has been given to the relevance of reference genome selection in variant calling (Thorburn et al. 2023; Rick et al. 2024; Akopyan et al. 2025). Using a combination of simulated and empirical data, Rick et al. (2024) demonstrated that different thresholds in the filtering of non-reference alleles influence the outcome of phylogenomic analyses. Importantly, they also showed that reference genome choice may indeed introduce biases that cascade through downstream analyses when variant calling is not accurate. Ultimately, this inaccuracy in variant discovery also affects the reliability and reproducibility of phylogenomic reconstructions. Based on these observations, Rick et al. (2024) concluded that choosing the reference genome for sequence alignment and variant calling is an important decision with implications for downstream analyses. However, apart from this general statement, their study falls short of providing practical solutions for the bioinformatic influences that need to be made to assemble sequences for phylogenomic datasets.

As for the influence of dataset characteristics, we note that the impact of any pre-existing bias arising from pre-existing nucleotide differences between the reference and the sample sequences is exacerbated in datasets characterized by comparatively low read coverage and incomplete marker representation. Under these conditions, the limited number of aligned short reads reduces the confidence of variant calling algorithms to distinguish between true nucleotide substitutions and sequencing errors. Both conditions are hallmarks that many phylogenomic studies of non-model organisms have in common (e.g., Zhang et al. 2019; Prasad et al. 2022; Thorburn et al. 2023).

Our experiments utilize the same software pipeline to rule out any influence software choice may have. Moreover, we included only exons that were consistently captured under each chosen reference and treated all alignment gaps as missing data when calculating genetic distances to ensure that the observed variation in p-distances between sequences from different runs is not an artifact of inconsistent sequence coverage. Based on these premises, the documented differences in variant calling outcomes must be caused by reference choice as the only varying parameter.

This leads us to the following conclusions: (1) Failure or error in aligning short reads against the reference are not the only mechanisms contributing to the error in variant calling (e.g., our short read alignment aligned by non-ingroup reference). (2) variant callers mistake real sequence differences with sequencer errors when a phylogenetically distant reference is used. This causes false negatives rather than false positives (e.g. Fig. 5a-b). (3) The reference sequence should be nearly identical (i.e., conspecific) with the sample sequence to minimize reference-related errors in variant calling (e.g. Fig. 5c). In the following section, we show how constructing a reference that meets these requirements can be achieved in cases where a conspecific reference genome is unavailable.

### Accurate outcomes through self-referencing

Variant calling offers clear advantages for genome-scale phylogenomic studies, as it mitigates biases and maximizes the use of available read information. However, none of the available phylogenomic pipelines for target-enrichment sequencing (Assexon, PHYLUCE, HybPiper) incorporate mechanisms to detect and correct errors arising from the use of phylogenetically divergent reference genomes. This limitation represents a critical methodological shortcoming that can compromise the accuracy of phylogenomic inferences.

Alternative approaches, which avoid variant calling altogether, such as *de novo* or BLAST-based assemblies, typically collapse all reads into a single consensus sequence (Johnson et al. 2016; Faircloth 2016). In this process information on polymorphisms is discarded. However, this information may be useful in phylogenetics. Moreover, it is relevant in making better variant calling decisions and to rule out sequencing errors. Without explicit evaluation of variants, these approaches cannot consider factors in the consensus making, such as read quality, or accommodate the models used in variant calling. This increases the likelihood of mis-assemblies and erroneous calls (Nielsen et al. 2011). In addition, these methods are prone to producing truncated contigs because regions of lesser similarity with probes or baits, particularly at the flanks of loci, are often trimmed for low read coverage.

To deal with the difficulties of reference choice in phylogenomics appropriately requires a tailored methodological adaptation of the bioinformatic pipeline and the analytical strategies (Fonseca et al. 2016). While we found that using the combined ingroup or an individual ingroup as reference in variant calling outperforms the use of a non-ingroup reference, we demonstrate that the self-referencing method produces the most accurate results both in terms of variant detection and phylogenetic consistency. The fact that variant callers were originally developed to deal with sequences from a single species that vary at a population level, probably explains why using sample-specific self-referencing is the best methodological pathway for accurately calling nucleotide variants even in a phylogenetically deeply structured sequence dataset. By minimizing biases introduced by external reference genomes, the self-referencing method provides a more accurate representation of genetic diversity. Based on these findings, we recommend updating assembly pipelines to generate a self-reference genome for each taxon prior to variant calling (Fig. 1). RGBEPP implements this strategy by performing a de novo assembly for each taxon as its own reference, allowing internally consistent variant discovery. By avoiding mapping to a distant genome, this method reduces false negatives, prevents systematic distortion of allele frequencies, and maintains the validity of downstream phylogenomic assumptions.

A sample specific self-reference is created in two simple steps. First, we perform a de novo assembly using the short sequencer reads into contigs. In target-enrichment studies, most of these contigs will represent the targeted loci. Second, we use BLASTx method to identify the ortholog that these contigs represent in reference to an available reference genome. Undertaking this filtering step also allows to exclusion any reads that are not on target. Performing it at the amino acid level allows the use of a phylogenetically more distant reference as amino acid sequences are more evolutionarily conserved than nucleotide sequences. After creation of this sample specific reference, raw reads are re-mapped to the nucleotide reference in order to extend the contigs and variant calling is performed to generate the final assembled contigs.

## Material and Methods

### Data Characteristics

We utilize the published exon capture dataset of Teasdale et al. (2016), which includes nuclear DNA sequences from 22 Australian species of land snails of the family Camaenidae. These sequences are moderately divergent as this group is of possible Miocene origin (Hugall and Stanisic 2011). This sequence dataset encompasses 2,648 exons from 490 orthologous protein-coding genes with an average exon length of 1,190 bp (228 to 6,261bp). Sequencing coverage per exon ranged from 41 to 235 (Teasdale et al. 2016). Each exon is a single-copy, orthologous locus as was qualified by Teasdale et al. (2016).

### Sequence Assembly and Alignment using RGBEPP

Data acquisition, preprocessing, ortholog identification, sequence alignment, and phylogenetic analyses were performed using the Reference Genome-Based Exon Phylogeny Pipeline (RGBEPP). This pipeline is written in the D programming language (Alexandrescu 2010) and is available on GitHub (https://github.com/starsareintherose/RGBEPP). It is also integrated in BioArchLinux (Zhang et al. 2025) under the GNU General Public License.

Following quality control and preprocessing of raw reads using fastp (Chen et al. 2018), we used two workflows that differed in reference implementation.

In the ‘RGBEPP_refmix’ workflow, the preprocessed reads are mapped to a reference genome using Bowtie2 (Langmead and Salzberg 2012). The resulting alignments are sorted, and duplicates are marked using SAMtools (Danecek et al. 2021). Variants are called and consensus sequences are generated with BCFtools (Danecek et al. 2021). Contigs are aligned based on codons using MACSE (Ranwez et al. 2018) and trimmed based on codon boundaries using trimAl (Capella-Gutiérrez et al. 2009) and a custom tool (delstop) written in D (Fig. 1).

In the ‘RGBEPP workflow’, instead of providing an external reference, we create a sample-specific reference by performing a *de novo* assembly using SPAdes (Prjibelski et al. 2020) and aligning these assemblies to the amino acid sequences of a global reference using DIAMOND BLASTx (Buchfink et al. 2021) and a custom tool, sortdiamond, to extract the orthologs that become sample-specific reference in variant calling. This reference is used in Bowtie2, SAMtools and BCFtools and is analogous to the ‘RGBEPP_refmix’ workflow described above (Fig. 1).

The configuration of all parameters, including customization options, and information on the input and output files of each step are summarized in Supplement 1.

### Variant Calling Experiments

To examine the influence of reference genome choice on variant calling, we trialed different references or reference types in 25 runs. In each run, we assembled the sequence dataset from scratch using the above workflow (Fig. 1) with different reference exomes for variant calling. In run 1, we used a non-ingroup representative (*Lottia gigantea*) as reference; in runs 2-23, we used a single ingroup representative each as reference; in run 24, we combined all 22 ingroup representatives in a composite reference; in run 25, we used sample-specific self-referencing. Runs 1-24 were performed using the RGBEPP_refmix workflow, run 25 was performed using the RGBEPP workflow (Fig. 1). To ensure that there were no false positive variant calls in run 25, we manually checked the short-read alignment by visualizing the mapped reads and called variants using the program IGV (Robinson et al. 2011).

All analyses were based on 1,004 exons from 399 orthologous protein-coding genes that were consistently represented in all partial reference exomes. The exome dataset for each reference genome was concatenated using catsequences (Creevey 2023).

### Assessing the Impact of Reference Choices

Uncorrected pairwise genetic distances between samples (p-distances) are used to quantify the differences in the proportions of detected variants. Distances were calculated using the ‘blastn’ method of the DistanceCalculator implemented in BioPython (Talevich et al. 2012).

First, we calculated distances between all samples in each MSA produced by the 25 runs and visualized the differences in heatmaps for each MSA. Secondly, we calculated distances between sequences of the same samples assembled under different reference choices. We applied Multidimensional Scaling (MDS) in scikit-learn (Pedregosa et al. 2011) to visualize these differences among the 22 samples in a two-dimensional scatter plot.

We performed a Maximum Likelihood tree reconstruction for each MSA produced by the 25 runs using RAxML-NG (Kozlov et al. 2019). In each case, we used ModelFinder (Kalyaanamoorthy et al. 2017) to select the best substitution model for these analyses.

We used 10 Maximum Parsimony plus 10 random starting trees and performed Tree Bisection and Re-connection to search for the best Maximum Likelihood tree with 1,000 bootstrap iterations. We calculated relative Robinson-Foulds (RF) distances (Robinson and Foulds 1980) with ETE3 (Huerta-Cepas et al. 2016) to quantify topological distances between phylogenies inferred from the MSAs produced under different reference selections. These relative RF distances are visualized as heatmaps. Additionally, we applied MDS to the relative RF distances to represent the relationships among phylogenetic trees in two-dimensional scatter plots.

## Funding Acknowledgment

This work was supported by Australian Biological Resources Study grant number RF213–12 to FK.

## Data Availability Statement

Software is available from GitHub: https://github.com/starsareintherose/RGBEPP. Data available from the Figshare [Once published, it will be made public].

## Author contribution

Guoyi Zhang: Conceptualization, Data curation, Formal analysis, Methodology, Resources, Software, Visualization, Writing – original draft, Writing – review and editing; Gerry Cassis: Writing – review and editing; Frank Köhler: Funding acquisition, Writing – original draft, Writing – review and editing.

## Acknowledgements

We gratefully acknowledge the constructive discussions with Luisa C. Teasdale (Centre for AgriBioscience, Australia), Adnan Moussalli (Museum Victoria, Australia), and Qi Feng (Capital Normal University, China). Our gratitude further goes to the dlang community for their insightful coding suggestions and to UNSW for providing the computational resources for GZ.

---

**Fig. S1.**
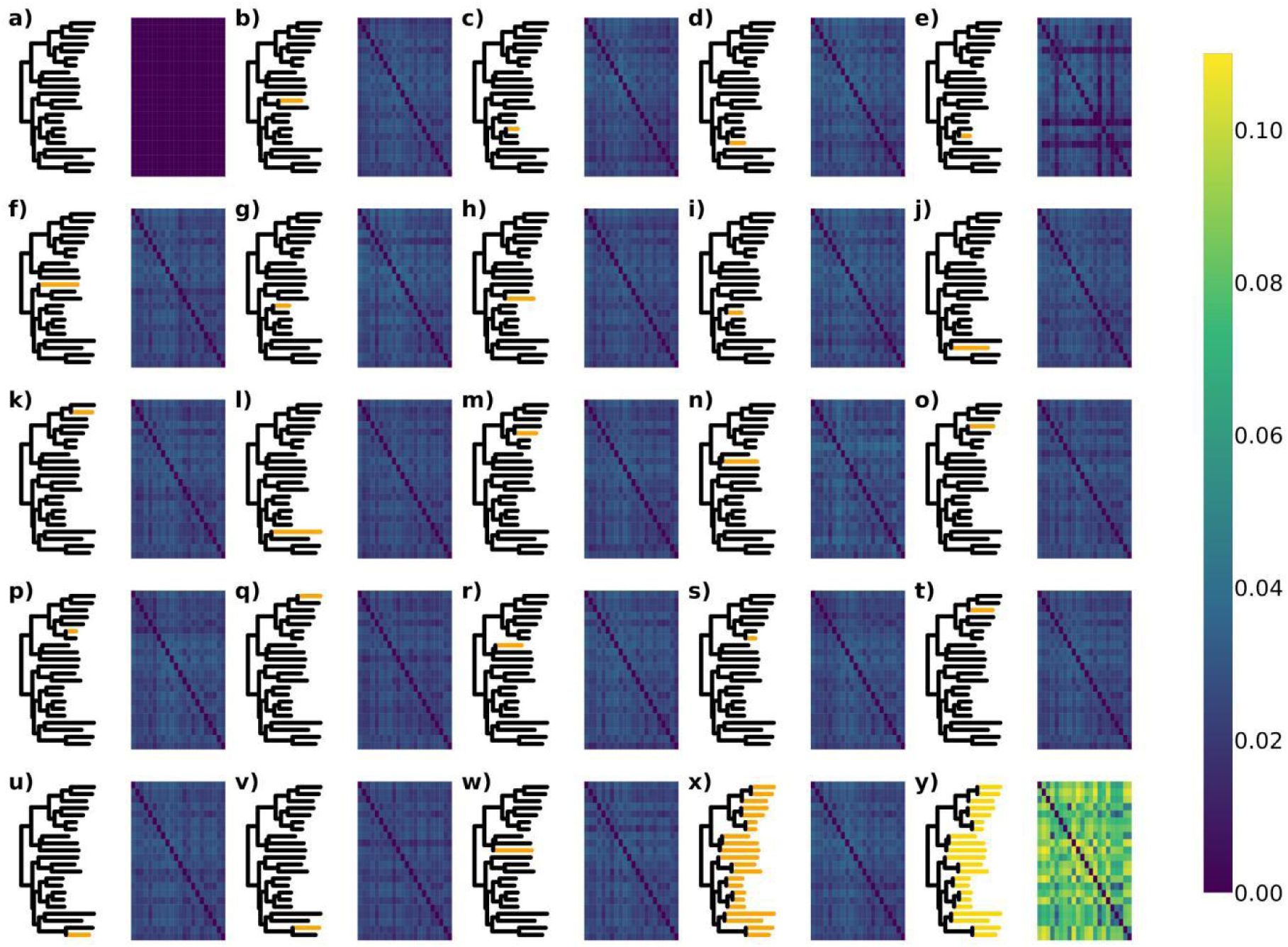
Heatmaps of pairwise genetic distances among the partial exomes in each of the MSA’s built using different reference exomes. a) non-ingroup reference (run 1); b-w) individual ingroup references (runs 2-23), referenced taxon highlighted in corresponding tree; x) combined ingroup reference (run 24); y) sample-specific self-reference (run 25).

**Fig. S2.**
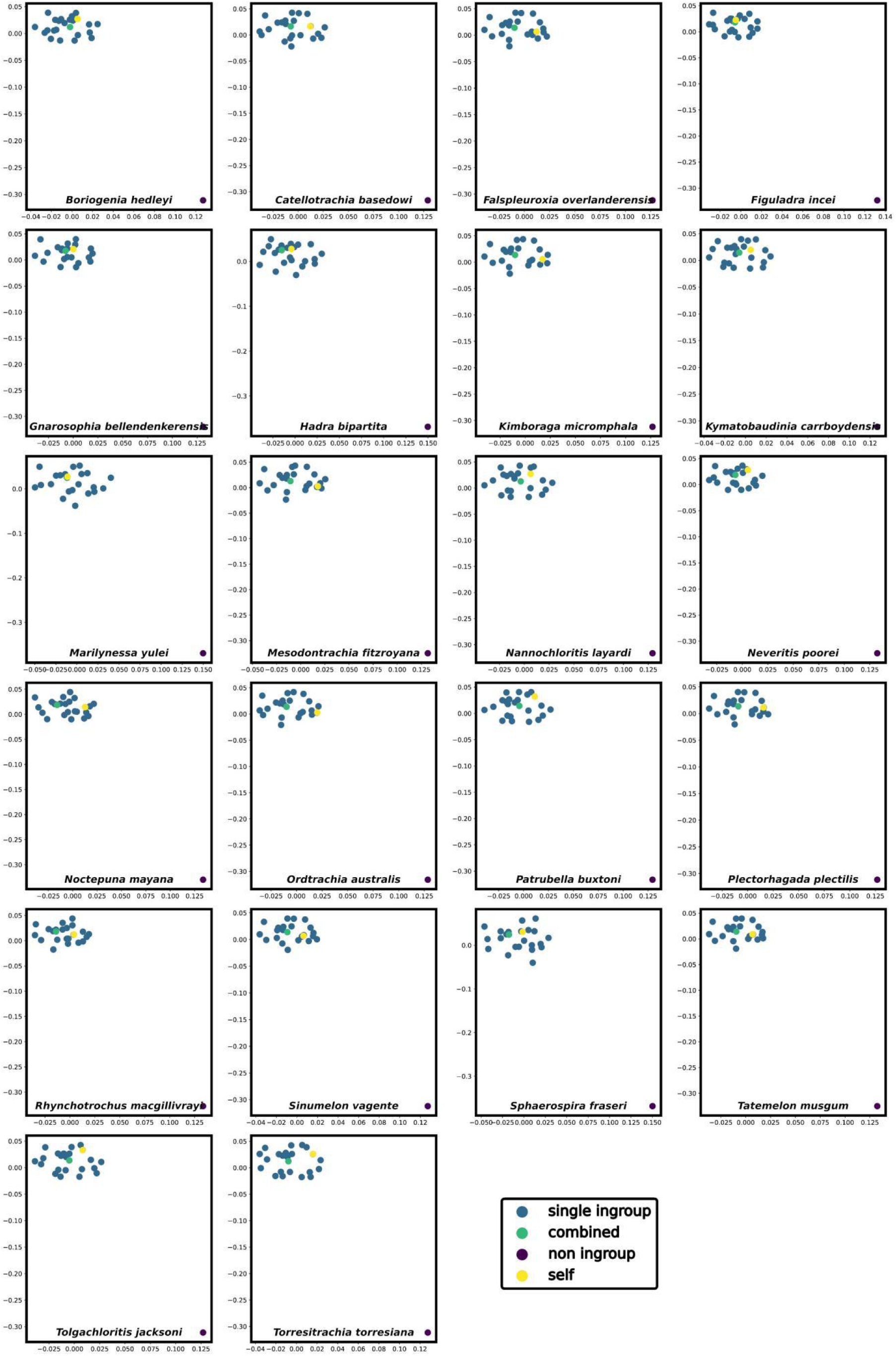
Scatter plots of multidimensional scaling (MDS) summarizing p-distances among ingroup taxa for each MSA. Colours indicate reference used for variant calling.

